# Low-Density Granulocytes are a novel immunopathological feature in both Multiple Sclerosis and Neuromyelitis optica spectrum disorder

**DOI:** 10.1101/668160

**Authors:** Lennard Ostendorf, Ronja Mothes, Sofie van Koppen, Randall L. Lindquist, Judith Bellmann-Strobl, Susanna Asseyer, Klemens Ruprecht, Tobias Alexander, Raluca A. Niesner, Anja E. Hauser, Friedemann Paul, Helena Radbruch

**Affiliations:** Department of Neuropathology, Charité-Universitätsmedizin Berlin, Corporate Member of Freie Universität Berlin, Humboldt-Universität zu Berlin, and Berlin Institute of Health; Department of Rheumatology and Clinical Immunology, Charité-Universitätsmedizin Berlin, Corporate Member of Freie Universität Berlin, Humboldt-Universität zu Berlin, and Berlin Institute of Health; Immunodynamics, Deutsches Rheuma-Forschungszentrum Berlin, A Leibniz Institute; Biophysical Analysis, Deutsches Rheuma-Forschungszentrum Berlin, A Leibniz Institute; Department of Nuclear Medicine, Charité-Universitätsmedizin Berlin, Corporate Member of Freie Universität Berlin, Humboldt-Universität zu Berlin, and Berlin Institute of Health; NeuroCure Clinical Research Center, Charité-Universitätsmedizin Berlin, Corporate Member of Freie Universität Berlin, Humboldt-Universität zu Berlin, and Berlin Institute of Health; Department of Neurology, Charité-Universitätsmedizin Berlin, Corporate Member of Freie Universität Berlin, Humboldt-Universität zu Berlin, and Berlin Institute of Health; Experimental and Clinical Research Center, Max Delbrück Center for Molecular Medicine & Charité –Universitätsmedizin Berlin, corporate member of Freie Universität Berlin, Humboldt – Universität zu Berlin, and Berlin Institute of Health, Germany; Department of Clinical Neurosciences, John Radcliffe Hospital, Oxford, UK; Clinical and Experimental Multiple Sclerosis Research Center, Charité-Universitätsmedizin Berlin, Corporate Member of Freie Universität Berlin, Humboldt-Universität zu Berlin, and Berlin Institute of Health; Fachbereich Veterinärmedizin, Institute of Veterinary Physiology, Freie Universität Berlin

## Abstract

**Objective:** To investigate whether low-density granulocytes (LDGs) are a immunophenotypic feature of patients with multiple sclerosis (MS) and/or neuromyelitis optica spectrum disorder (NMOSD).

**Methods:** Blood samples were collected from 26 patients with NMOSD and 20 patients with MS, as well as from 18 patients with Systemic Lupus Erythematosus (SLE) and 23 Healthy Donors (HD). We isolated peripheral blood mononuclear cells (PBMCs) with density gradient separation and stained the cells with antibodies against CD14, CD15, CD16, and CD45, and analysed the cells by flow cytometry or imaging flow cytometry. We defined LDGs as CD14^-^CD15^high^ and calculated their share in total PBMC leukocytes (CD45^+^) as well as the share of CD16^hi^ LDGs. Clinical data on disease course, medication, and antibody status were obtained.

**Results:** LDGs were significantly more common in MS and NMOSD than in HDs, comparable to SLE samples (median values HD 0.2%, MS 0.9%, NMOSD 2.1%, SLE 4.3%). 0/23 of the HDs, but 17/20 NMOSD and 11/17 MS samples as well as 13/15 SLE samples had at least 0.7 % LDGs. NMOSD patients without continuous immunosuppressive treatment had significantly more LDGs compared to their treated counterparts. LDG nuclear morphology ranged from segmented to rounded, suggesting a heterogeneity within the group.

**Conclusion:** LDGs are a feature of the immunophenotype in some patients with MS and NMOSD.

## Background

Low-Density Granulocytes (LDGs) are neutrophilic granulocytes that remain in the fraction of peripheral blood mononuclear cells (PBMC) after density gradient separation. When first described, they were noted to be a feature of rheumatologic diseases such as Systemic Lupus Erythematosus (SLE) (1), while more recent works described LDGs in various conditions such as asthma (2), tuberculosis (3), and psoriasis (4). LDGs have been implicated in the pathogenesis of SLE by producing type I interferons and undergoing spontaneous NETosis (generation of Neutrophil extracellular traps), thus providing potentially immunogenic nuclear matter (5,6). Their role in the pathogenesis of other diseases is less clear.

The origin of the LDGs is not yet explained and it has been proposed that they represent immature or degranulated neutrophils. Interestingly, LDGs in SLE display an immature nuclear structure even though they express the surface markers of mature granulocytes, such as CD16. (5,6)

Neuromyelitis Optica Spectrum Disorder (NMOSD) and Multiple Sclerosis (MS) are neuroinflammatory conditions of unknown etiology. The evidence for the involvement of granulocytes in the pathogenesis of MS has been scarce so far, while some early studies indicate a role of neutrophils in NMOSD pathogenesis. (7,8)

In this study, we provide the first evidence for the existence of LDGs in MS and NMOSD and thus add another puzzle piece to the role of the innate immune system in the pathogenesis of these diseases.

## Methods

### Patient selection

SLE patients were recruited from the rheumatologic outpatient departments at Charité Universitätsmedizin Berlin. Data on MS- and NMOSD-patients were derived from ongoing observational studies at the NeuroCure Clinical Research Center, Charité – Universitätsmedizin Berlin. These studies were approved by the ethics committee of the Charité – Universitätsmedizin Berlin (MS: EA1/163/12; NMOSD: EA1/041/14) and conducted according to the 1964 Declaration of Helsinki in its currently applicable version. Written, informed consent was obtained from all participants in the study.

MS-diagnosis was made according to the 2010 revised McDonald criteria, NMOSD-diagnosis was made according to the international consensus diagnostic criteria for NMOSD 2015.

All MS and NMOSD patients were a stable phase of disease with a minimum of three months since the last relapse and last glucocorticoid pulse. We excluded patients if they received the diagnosis of an additional autoimmune, infectious or malignant condition. We correlated the LDG percentage with clinical information such as Expanded Disability Status Scale (EDSS) score, number of relapses, treatment, and duration of disease as well as titres for anti-MOG and anti-AQP4 antibodies.

### PBMS isolation, Staining and Flow Cytometry

We isolated the PBMC layer with density-gradient separation of 10ml heparin-anticoagulated blood, using the Ficoll-Paque PLUS gradient (GE Healthcare). PBMCs were washed once in phosphate-buffered saline (PBS) supplemented with 3% bovine serum albumin (BSA) and subsequently stained with the following antibody-conjugates: CD45 FITC (clone 5B1, Miltenyi), CD14 Cy5 (clone TM1, DRFZ), CD15 PE-Cy7 (clone W6D3, Biolegend), CD16 PE (clone 3G8, Biolegend), CD14 APC-Cy7 (clone 63D3, Biolegend) and CD11c Pacific Blue (clone M1/70, Biolegend). We acquired the sample using a FACSCanto (BD Biosciences) or MACSQuant (Miltenyi) cytometer. For imaging flow cytometry (Amnis), we isolated and stained the cells as described above, fixed the sample with the Foxp3 / Transcription Factor Staining Buffer Set (eBioscience) and acquired the cells on an Amnis ImageStream X Mark II. All cytometry experiments were performed in accordance with the “Guidelines for the use of flow cytometry and cell sorting in immunological studies”. (9)

### FACS Data Analysis

We excluded sub-cell-sized detritus and doublets based on scatter characteristics and gated on CD45^+^ leukocytes. Of these, we defined LDGs as CD14^-^CD15^high^ and calculated the ratio of LDGs of all CD45+ PBMCs (Fig 1a). We then determined the fraction of CD16^high^ LDGs. FACS Data Analysis was performed using FlowJo Version 10.4.1 (FlowJo LLC). We analyzed the ImageStream data with IDEAS Version 6.2 (Amnis).

**Figure 1.**
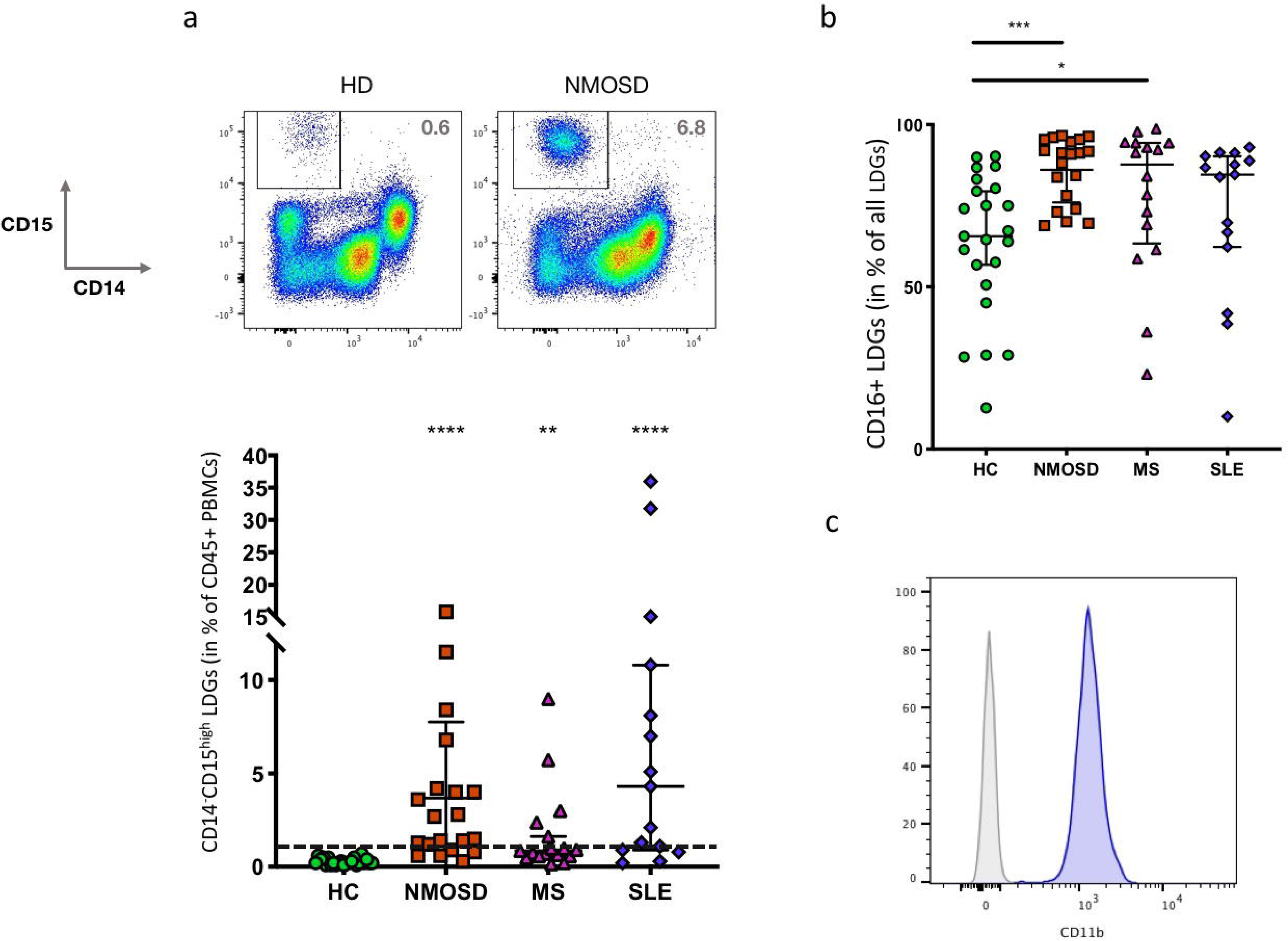
a) LDGs (CD14-CD15^high^) are a feature NMOSD and MS but not HDs (HD: n = 23, NMOSD: n = 20, MS: n = 17 SLE: n = 15). Bars indicate median and 95% Confidence Intervals. The horizontal bar indicates 0.7%, a cut-off that seperates all HDs from 17/20 NMOSD, 11/17 MS samples as well as 13/15 SLE. Kruskal-Wallis test with Dunn’s Correction for multiple testing **: p < 0.005, ****: p < 0.0001) b) Most LDGs are CD16^high^, but there are more CD16^high^ LDGs in NMO and MS than in HDs. There are some outliers with very low levels of CD16^high^ LDGs. Bars indicate median and 95% Confidence Intervals. Kruskal-Wallis test with Dunn’s Correction for multiple testing. (HD: n = 23, NMOSD: n = 20, MS: n = 16 SLE: n = 15), Test, *: p < 0.05, ***: p < 0.001) c) LDGs are CD11b+ (Gated on CD14-CD15+, gray: fluorescence minus one (FMO) control)

### Statistical Analysis

The percentage of LDGs of different disease states and the percentage of CD16^high^ LDGs were compared to HDs using the Kruskal-Wallis test with Dunn’s Correction for multiple testing. For the difference between two groups, we used the Mann-Whitney test. All statistical analysis was performed in GraphPad Prism Version 7.0e for Mac OS X (GraphPad Software).

## Results

LDGs are a feature of many inflammatory conditions and of unknown pathogenic significance. To elucidate whether LDGs also exist in NMOSD and MS, we performed FACS analyses of 76 peripheral blood samples.

### Cohort Description

We conducted the analysis of 17 MS patients as well as 20 patients with NMOSD. 15 patients with SLE and 23 HDs served as controls. Median EDSS scores were 4.0 in the MS and 3.0 in the NMOSD group. The most common treatments in the MS group were Dimethylfumarate (6/17) and beta Interferon (4/17). The median number of relapses in the relapse-remitting MS (RRMS) patients was 2.5. NMOSD Patients were most often treated with Rituximab (8/20), Mycophenolate (4/20) and Azathioprine (4/20). (Table 1)

**Table 1:**
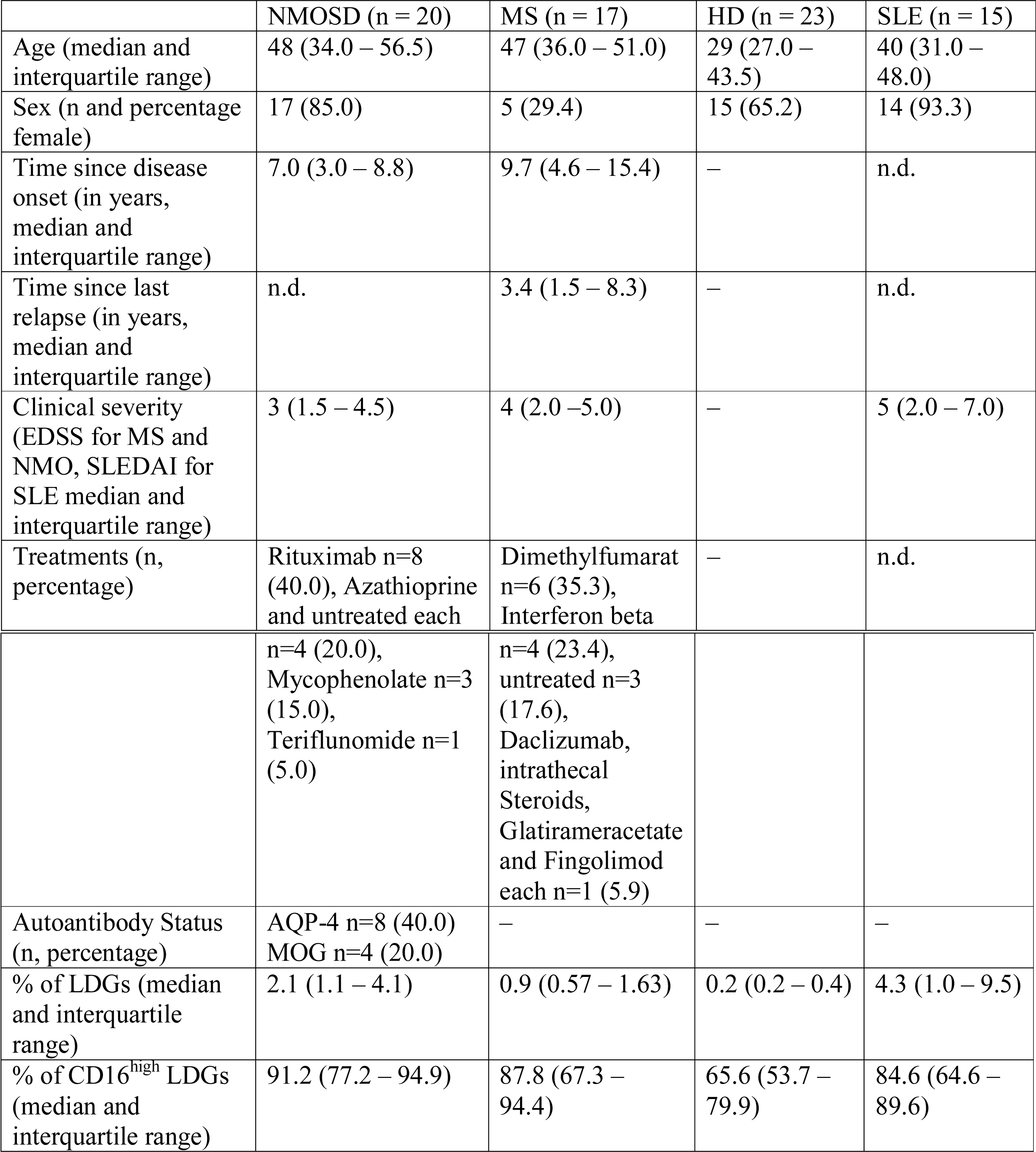
Epidemiological Information of the Study Population.

### Low-Density Granulocytes are a feature of both multiple sclerosis and neuromyelitis optica

In comparison to healthy controls, both MS and NMO patients feature a significantly higher percentage of LDGs. SLE samples displayed the highest frequencies of LDGs. (MS: 0.9%, NMO 2.1%, SLE: 4.3%, HD: 0.2%, median fraction of CD45^+^ cells) (Fig 1a). None of the HDs, but 17/20 NMOSD and 11/17 MS samples as well as 13/15 SLE samples had at least 0.7 % LDGs. The fraction of CD16^high^ LDGs differed both between and within groups (Fig 1b), but was significantly higher in NMOSD and MS compared to HCs. LDGs uniformly expressed CD11b (Fig 1c).

### LDGs show a heterogeneous nuclear morphology

We analyzed the nuclear morphology of the LDGs of a NMO patient. While some of the CD14^-^, CD15^high^ cells displayed the segmented nuclear shape typical of mature granulocytes, others showed band-formed or rounded nuclei, which are traditionally thought to be immature forms. (Fig 2)

**Figure 2:**
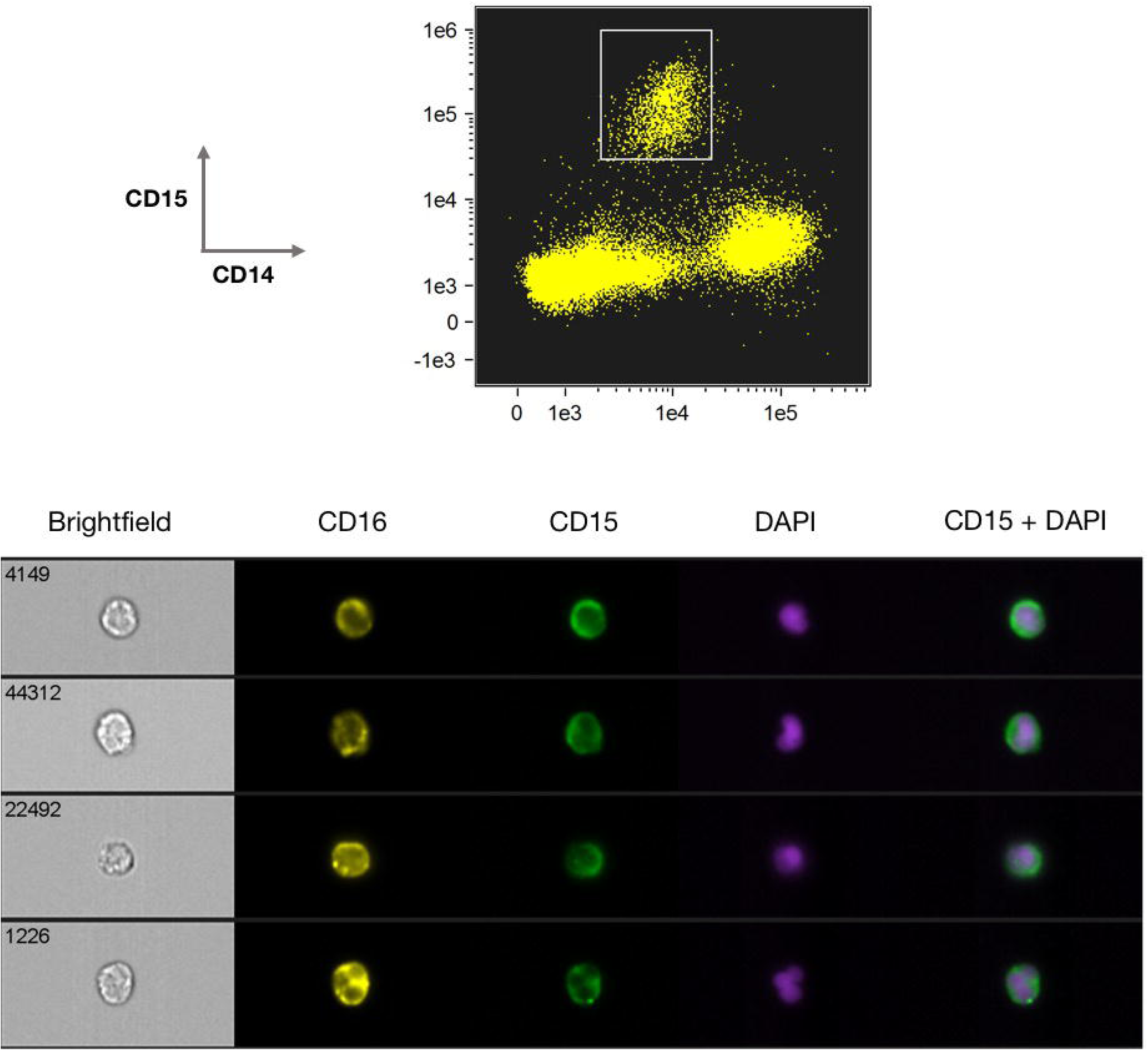
Imaging flow cytometry analysis of LDGs from a patient with NMO revealed heterogenous nuclear shapes, ranging from round to multisegmented.

### Untreated NMOSD patients show higher LDG fractions

We looked for associations between the fraction of LDGs in MS and NMO patients and their clinical characteristics such as time since first manifestation of disease, number of relapses, severity of disability as judged by the EDSS score, type of treatment and antibody status, but did not find any significant correlations. (Data not shown) NMOSD patients who did not receive a continuous immunosuppressive treatment had significantly higher LDG percentages compared to their peers receiving treatment. (median LDGs treated 1.4% untreated 7.9%, p < 0.01). The LDG fraction did not correlate with clinical severity.

## Discussion

This study demonstrates the presence of LDGs in MS and NMOSD. In contrast to MS, in NMO classical granulocytes are thought to play a pivotal role as they can be found in tissue biopsies (7) and alterations in neutrophil function have been described (8). We recently described CD11b+ leucocytes in the PBMCs of MS patients that were characterized by increased activation of NAD(P)H oxidase (NOX). (10) In these, NOX activation correlated with clinical activity. While we assumed these cells to be monocytes at the time, the presence of LDGs in the peripheral blood mononuclear cell (PBMC) fraction could pose an alternative explanation for our previous results.

We found the LDG fraction of NMOSD and MS patients to contain significantly more CD16^high^ cells, a marker indicating neutrophil maturity. As previously described for LDGs in SLE, LDG nuclei in NMOSD vary between mature segmented and immature round forms. In previous studies, the immature nuclear morphology was associated with a expression of surface markers characteristic of mature cells, suggesting that LDGs do not fit into the traditional maturity spectrum of granulocytes. (6)

This work adds MS and NMOSD to the diverse list of inflammatory conditions featuring LDGs. In our opinion, this makes it more likely that LDGs are an epiphenomenon of ongoing inflammation rather than a causal part of the pathogenesis of a specific disease, but further research is needed to establish the role of these cells in different conditions.

To establish the potential of LDGs as a biomarker in neuroinflammatory diseases or inflammation in general, analysis of larger and longitudinal cohorts should be performed to uncover potential correlations with disease severity or prognosis.

